# Initial Condition Decision to Ensure Reliable Circadian Phase Estimation with Shorter-Term Wearable Data

**DOI:** 10.1101/2025.11.01.686004

**Authors:** Dongju Lim, Juhyeon Kim, Jee Hyun Kim, Jae Kyoung Kim

## Abstract

Various studies in the medical field highlight the importance of circadian medicine, which optimizes treatment timing based on patients’ circadian phases. While the circadian phase has been measured using dim light melatonin onset (DLMO), the gold standard marker, its high cost and time-intensive nature have led to the development of alternative estimation methods. Among these, the most promising method is using ordinary differential equation (ODE) models, which simulate the circadian rhythm by using a light exposure profile estimated from sleep data. These ODE models require an initial condition (IC) representing the initial state of the circadian rhythm, which is unknown in real-world settings. However, it is unclear how the uncertainty of IC affects the accuracy of circadian phase estimation. In this study, by using sleep data collected from 28 shift workers using ActiWatch (mean duration = 56 days, range = 34–75 days), we found that ≥18 days of sleep data are required for the circadian phase to become independent of the subjective IC choice. The result showed that without an accurate IC, circadian phase estimation is dependent on subjective IC choice, meaning that circadian phase estimates in the first 17 days are not reliable. Indeed, these days were reduced to 13 – 15 days on average when previous studies’ IC estimation methods were used. To further shorten this length, we developed new IC estimation methods—period- and work history-based sleep methods—that incorporate daily variations in sleep history. Notably, the new methods reduced the number of days required for reliable circadian phase estimation to about seven days. Hence, our approach allows a larger portion of circadian phase estimates from given sleep data to be used as reliable information. The superiority of our methods paves the way for improved circadian phase estimation, ultimately enhancing the practicality of chronotherapy applications.

## Introduction

The circadian rhythm, a bio-oscillator with a 24-hour period, regulates various biological processes such as metabolism and homeostasis (Klein and Weller, 1970; Narasimamurthy et al., 2012; Black et al., 2019; Martchenko et al., 2020). Accordingly, the treatment of disease caused by an abnormal biological process is closely related to the circadian rhythm. For example, the treatment of cancers (Mormont and Levi, 2003; Hsu et al., 2016; Kim et al., 2023), cardiovascular diseases (Portaluppi and Lemmer, 2007; Stranges et al., 2015; Hermida et al., 2020; Wei and Diekman, 2025), and autoimmune diseases (To et al., 2009; To et al., 2011; Wang et al., 2018) has been shown to benefit from determining the optimal timing of drug administration in alignment with the circadian rhythm. Consequently, the importance of the treatment that considers an individual’s circadian phase, chronotherapy, has been highlighted (Kim et al., 2020). Effective chronotherapy requires accurate measurement of the circadian phase, which can vary largely depending on an individual’s lifestyle factors, such as sleep patterns (Paine and Gander, 2016). Conventionally, this has been achieved by measuring the dim light melatonin onset (DLMO), the gold standard biomarker for the circadian phase (Lewy et al., 1999; Benloucif et al., 2005; Pandi-Perumal et al., 2007).

However, DLMO measurement requires subjects to remain in a dim-light room for an extended period while saliva or blood samples are collected to measure melatonin levels, resulting in considerable time and cost burdens (Zeitzer et al., 2007; Glacet et al., 2023; Kennaway, 2023). To address these limitations, alternative computational methods for estimating DLMO have been developed (Reiter et al., 2020). Among them, the most promising approach is to use an ordinary differential equation (ODE) model based on the Van der Pol oscillator (Stone et al., 2020b). This model simulates core body temperature (CBT) rhythms using a light exposure profile as an input (Forger et al., 1999; Jewett et al., 1999; Hilaire et al., 2007). By subtracting seven hours from the minimum point of the simulated CBT (CBTmin), DLMO can be estimated (Hilaire et al., 2007; Woelders et al., 2017; Stone et al., 2019; St Hilaire et al., 2020; Stone et al., 2020a; Huang et al., 2021; Lim et al., 2025). While this method ideally requires an accurate light exposure profile, collecting such data remains challenging. One solution to this limitation is to estimate the light exposure profile from the sleep pattern data, assuming light exposure occurs only during wakefulness (Postnova et al., 2016; Hong et al., 2021; Knock et al., 2021; Song et al., 2023; Varma et al., 2023; Lim et al., 2024; Song et al., 2024; Lim et al., 2025; Song et al., 2025).

Nevertheless, a fundamental limitation remains in the ODE-based circadian phase estimation method: setting an accurate initial condition (IC). Accurately determining IC requires knowing the state of the circadian rhythm at the initial time point (i.e., the beginning of the sleep data). Such an initial state is usually unknown in real-world settings, as it primarily depends on the sleep pattern before the initial time point (i.e., sleep history), which is also unavailable. Consequently, to determine the IC, previous studies estimated the sleep history with their own approaches (Stone et al., 2019; Cheng et al., 2021; Huang et al., 2021; Mayer et al., 2023; Song et al., 2024). For instance, Huang et al. estimated sleep history using a constant sleep pattern with fixed timing: 23:30 – 7:30 (Huang et al., 2021). As this method does not account for inter-individual variation (e.g., duration, timing), Stone et al. developed an alternative approach that partially considers inter-individual variation by adjusting sleep timing based on the median phase of individuals’ sleep (Stone et al., 2019). However, this approach still assumes the same sleep patterns every day, neglecting intra-individual variation (i.e., daily variation).

In this study, by using sleep data collected from 28 shift workers via ActiWatch (mean length = 56 days, range = 34-75 days), we found that simulated circadian phases become independent of the arbitrary choice of IC after at least 18 days of simulations. This implies that the simulated circadian phase during the first 17 days cannot be reliably used without accurate IC estimation. Indeed, when previous sleep history estimation approaches were used to estimate IC, the required simulation days were reduced to 13 – 15 days on average. To further shorten this, by accounting for intra-individual variation in sleep, we developed new approaches: period- and work history-based sleep methods. With our new approaches, the number of days required for reliable circadian phase estimation is reduced to just about 11 and 7 days in the period- and work history-based sleep methods, respectively. We expect the superior performance of our IC estimation methods to provide a milestone in improving the accuracy of circadian phase estimation.

## Methods

### Data collection

We utilized the data collected by Lee et al. (Lee et al., 2024), which included sleep-wake patterns and work schedules from 28 female nurses. Sleep-wake data were recorded using Actiwatch (Actiwatch2 or Actiwatch Spectrum Pro), and their work shifts fell into four categories: off (no work), day (7:00-15:00), evening (15:00-23:00), and night (23:00-7:00), with each shift lasting up to four days. Participants had no physiological and neural disorders, and their age ranged from 23-52 (mean: 28.66±2.97 years) (Lee et al., 2024). Further details on data collection are available in Lee et al. (Lee et al., 2024).

### Model simulation

To simulate the circadian rhythm of shift workers, we utilized mathematical models. These were ordinary differential equation (ODE) models based on Van der Pol oscillators: simpler (Forger et al., 1999), higher-order (Jewett et al., 1999), and nonphotic models (Hilaire et al., 2007). Given that models’ input is light exposure profiles, we estimated the light exposure profile from the sleep data by assuming 250lux during wakefulness and 0lux during sleep, following previous studies (Postnova et al., 2016; Hong et al., 2021; Knock et al., 2021; Song et al., 2023; Varma et al., 2023; Lim et al., 2024; Song et al., 2024; Lim et al., 2025; Song et al., 2025). The estimated light exposure profile was used as an input for mathematical models to simulate the CBT rhythm (Fig. 1a). From this simulated CBT rhythm, we identified the CBTmin using the ‘findpeak’ function in MATLAB 2023b.

**Fig. 1.**
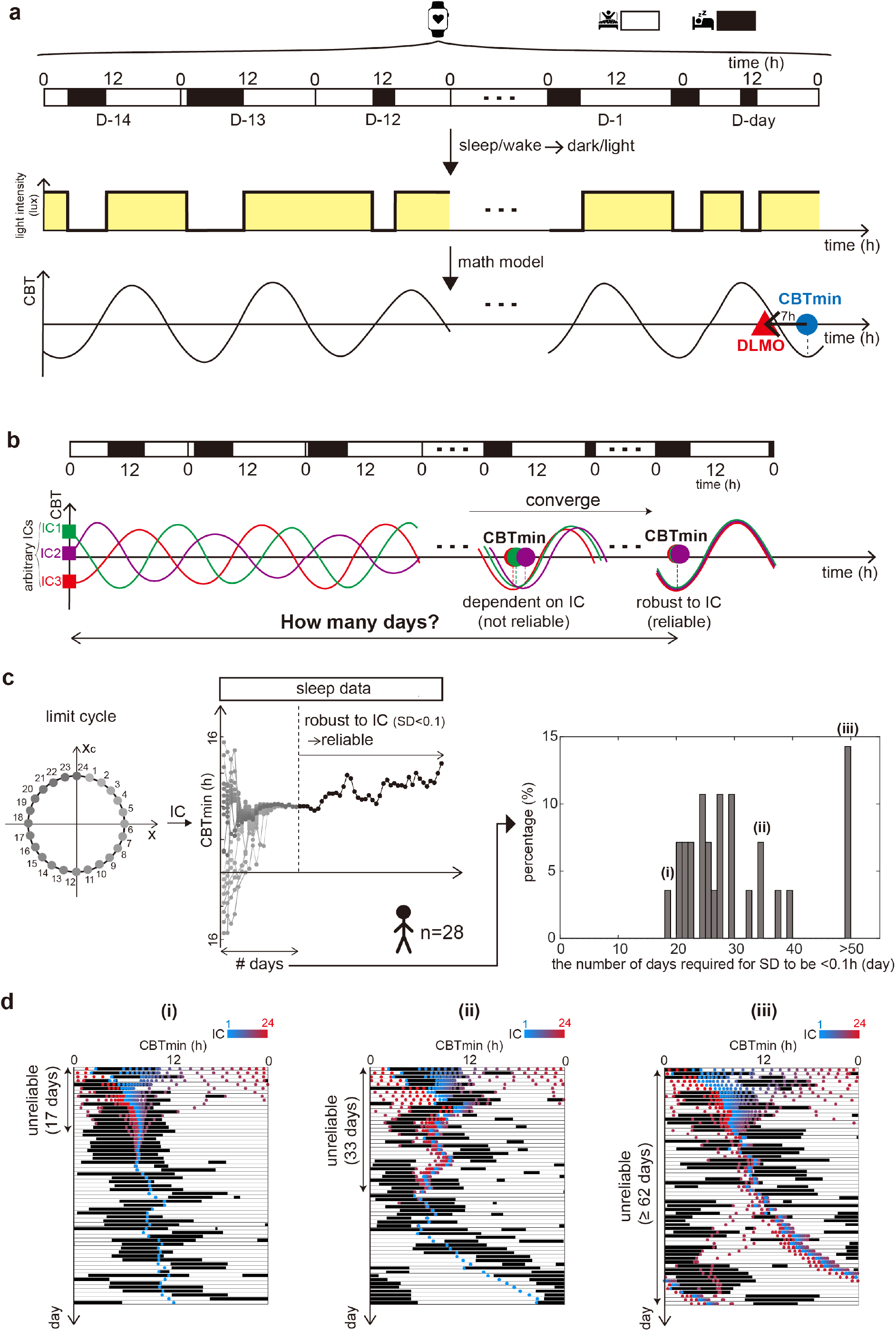
Estimation of circadian phase with mathematical model simulations robust to initial condition requires prolonged sleep-wake data. (a) The sleep/wake pattern data from wearable devices can be used to estimate light exposure, assuming exposure to light during waking hours and darkness during sleep. This light profile serves as an input for a mathematical model that simulates the circadian rhythm of the core body temperature (CBT). This allows us to identify the minimum point of the CBT (CBTmin, blue circle) on a specific day (D-day) and predict dim light melatonin onset (DLMO, red triangle) by subtracting seven hours from the CBTmin. (b) While the simulated circadian phases are heavily dependent on initial conditions during the early stage of the simulation, they eventually converge to the same phases regardless of initial conditions (i.e., become robust to IC), enabling reliable circadian phase estimation. However, the length of sleep-wake data required to ensure estimation that is robust to the initial condition is not known. (c) We investigated the minimal length of sleep data required for convergence across 24 different initial conditions (ICs; circles on the left) with each representing evenly distributed points on a limit cycle derived from a regular sleep pattern (22:00 - 6:00; See Methods for more details). Specifically, using the sleep/wake pattern data of 28 shift workers, we calculated the number of days required for the standard deviation (SD) of CBTmin values from 24 ICs to drop below 0.1h. Results show that at least 18 days are required for all tested subjects for reliable circadian phase estimation. Furthermore, there are also some cases that provide no reliable circadian phase estimation even when their entire sleep data (>50 days) is used in simulation. (d) The figure of the sleep pattern (black regions) and corresponding CBTmin (colored dots) for three subjects who required 18 days (i), 34 days (ii), and >63 days (iii) for reliable circadian phase estimation. Longer-term data is required for reliable estimation as sleep patterns become more irregular.

### Calculation of CBTmin robust to the initial condition

To investigate the influence of initial conditions on CBTmin calculation, we generated 24 different initial conditions and analyzed when the corresponding trajectories converged. First, we simulated the circadian rhythm entrained to a constant sleep pattern (22:00 – 06:00) for 50 days, starting from the initial condition (*x, x*_*c*_, *n*) = (0.5, −0.25, 5). From this entrained rhythm, we selected 24 hourly time points (1:00, 2:00, …, 24:00), using them as 24 initial conditions (points on the left circle in Fig. 1c). Using these 24 initial conditions, we calculated 24 CBTmin trajectories with real sleep-wake pattern data collected from shift workers (middle plot in Fig. 2c). We then identified the first day when the standard deviation of 24 CBTmin values, excluding outliers (i.e., greater than Q3+1.5 IQR or less than Q1-1.5 IQR, where Q1 and Q3 are the 1^st^ and 3^rd^ quantiles, respectively, and IQR is Q3-Q1) (Ranga Suri et al., 2019), fell below 0.1h. At and after the identified day, the mean of 24 CBTmin values (excluding outliers) was used as an IC-robust CBTmin in our analysis (black points in the middle plot in Fig. 2c). This process was repeated for the simpler, higher-order, and nonphotic models. In subjects with messy sleep patterns, the convergence of CBTmin values from distinct IC was slower, so in some cases, CBTmin did not converge even when using the entire sleep data (Fig. 1d (iii)). As such subjects couldn’t provide an IC-robust CBTmin, they were excluded from further analysis: three subjects for each of the three models.

**Fig. 2.**
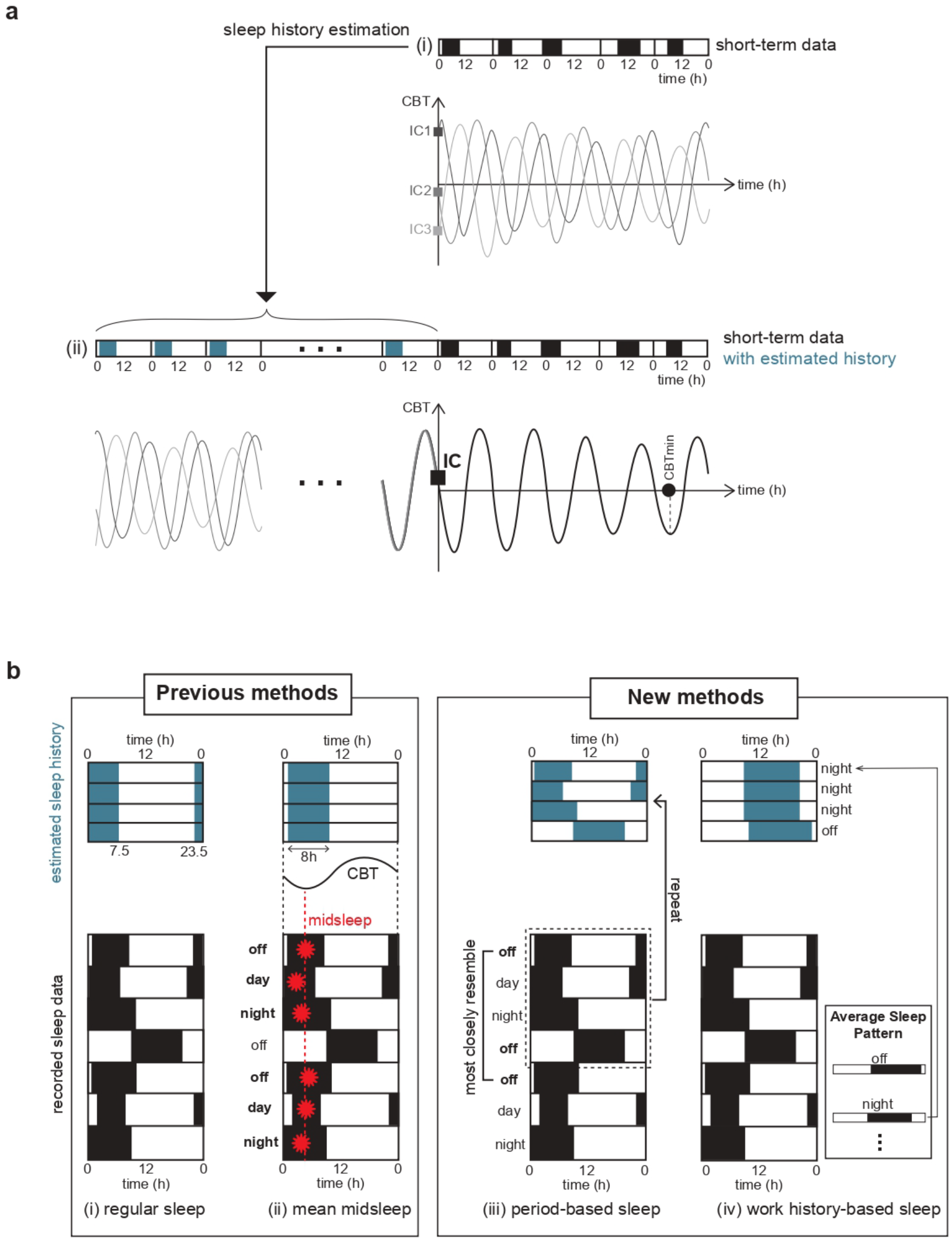
The methods for estimating sleep history to lengthen sleep-wake data. (a) When only short-term sleep data is available in real-world situations (i), the estimation of the circadian phase using the mathematical model heavily depends on the subjective choice of initial conditions (ICs). Therefore, selecting an accurate IC is crucial for accurate circadian phase estimation, and IC is determined by the circadian state at the initial time point (i.e., the beginning of the sleep data). To estimate this, previous studies have estimated sleep history (blue-shaded region in (ii)) and determined the IC by simulating the mathematical model for this history. Then, using this IC, we can simulate circadian phases, reducing the effect of subjective IC choice from the recorded short-term sleep data (black regions). (b) Previously, two classical methods have been used to estimate unknown sleep history: the regular sleep and mean midsleep methods (left). The regular sleep method estimates an unknown sleep-wake history as a constant 8-hour-length sleep starting at a fixed time (e.g., 23:30) (i). The mean midsleep method also assumes a constant 8-hour-length sleep at a fixed time while reflecting the actual sleep data. This method assumes sleep-wake history as a constant sleep, where historical CBTmin equals the average of midsleeps (red dotted line, ii) derived from only the midsleep values following non-night shifts (bold, ii) in recorded sleep data. However, as these methods assume constant sleep-wake patterns without intra-individual variation (i.e., daily variation) in sleep, we propose two new methods: the period-based sleep method and the work history-based sleep method (right). The period-based sleep method estimates sleep-wake history by assuming that a specific period in the recorded sleep data is repeated in the sleep history. The starting point of the period is fixed on the first day, and the endpoint is identified to ensure a seamless connection with the subsequent period. Specifically, among all days that share a common work pattern with the first day (bold, iii), the day whose sleep pattern most closely resembles that of the first day is selected. The day preceding this selected day is chosen as the endpoint of the period (dotted box, iii). This period is then repeated to create a sufficiently long sleep pattern, ensuring robustness to IC. The work history-based sleep method estimates sleep history by leveraging known work history. Specifically, the average sleep pattern corresponding to each work pattern is calculated (box, iv), and this average sleep pattern is matched to the work history (black arrow, iv). Then the matched pattern is repeated to generate a sufficiently long sleep pattern, ensuring robustness to IC.

### Implementation of previous IC estimation methods

In this study, we employed two classical sleep history estimation methods: the regular sleep method (Fig. 2b (i)) and the mean midsleep method (Fig. 2b (ii)). For the regular sleep method, we assumed a constant sleep pattern (sleep during 23:30 – 07:30) for 50 days, following a previous study (Huang et al., 2021). For the mean midsleep method, we assumed an 8-hour constant sleep pattern in the sleep history, but the sleep timing was determined by the mean midsleep, the average of median timing of sleep (i.e., midsleep), following a previous study (Stone et al., 2019). Specifically, we first calculated the daily midsleep phase across all days in the given sleep data and then determined their mean value (i.e., mean midsleep). For the mean midsleep calculation, following a previous study (Stone et al., 2019), we averaged the midsleep only on days following a non-night shift (indicated in bold in Fig. 2b (ii)). We then generated a 50-day sleep pattern with a constant 8h sleep pattern whose CBTmin was equal to the mean midsleep values and assumed it as the sleep history.

### Implementation of new IC estimation methods

Our new methods, the period-based sleep method (Fig. 2b (iii)) and the work-history-based sleep method (Fig. 2b (iv)), utilize not only sleep patterns but also work pattern data under the assumption that the sleep timing is closely linked to work schedules. Originally, work schedules were categorized into four types: off, day, evening, night. However, a more detailed classification was necessary to accurately link sleep patterns with work schedules. For example, even for the same night shift, sleep timing may differ depending on the previous day’s schedule; if the previous day was off, sleep could occur at any time between 00:00 and 23:00, whereas if the previous day was also a night shift, sleep would only be possible between 07:00 and 23:00. To address this, we refined the work schedule classification into six categories: off after night, off after non-night, first night, non-first night, day, and evening. This refined work schedule was utilized to implement the period- and work history-based sleep methods.

The period-based sleep method identifies a repetition period and repeats it in the sleep history. Thus, the key step in this method is to determine the starting point and endpoint of the repetition period. The starting point was fixed as the first day of the recorded sleep data, while the endpoint was identified as the day before the most similar sleep pattern to the starting point (i.e., the first day) to seamlessly connect consecutive periods. To find the most similar day to the first day, we first identified all days with the same work pattern as the first day (indicated in bold in Fig. 2b (iii)). Among them, we chose the day whose sleep pattern was most similar to that of the first day. Specifically, the similarity in sleep pattern between two sleep patterns was evaluated by transforming these sleep patterns into sleep vectors, where each element (1 for sleep and 0 for wakefulness) represents a minute of the day (i.e., 1440 in length for a day) and calculating the number of indices at which both sleep vectors have the same entry (i.e., both sleep or wake) (Lim et al., 2025). If the chosen day was within the first four days, which is the maximum number of consecutive days with the same work pattern, we recalculated the most similar day only among the days after the fourth day to prevent a short repetition period, which could reduce the reflection of daily sleep variation in the sleep history. Additionally, if no day shared the same work pattern as the first day, we selected the day having the most similar sleep pattern to the first day among all days in the sleep data after the fourth day. The day preceding this chosen day was then selected as the endpoint of the repetition period (dotted box in Fig. 2b (iii)). Finally, this period was repeated in the sleep history until its length exceeded 300 days.

The work history-based sleep method constructs sleep history by leveraging historical work schedule data. Specifically, it matches work patterns and sleep patterns by calculating the average sleep pattern for each of the work patterns (box in Fig. 2b (iv)). To calculate the average sleep pattern, we first generated a sleep vector for each day as in the preceding paragraph. These sleep vectors were then grouped by their corresponding work pattern. For each work pattern, we computed the average vector of sleep vectors across all associated days. Then we smoothed this average vector by using the ‘smoothing’ function (window size = 80) in MATLAB 2023b. Next, we determined the threshold such that the proportion of entries in the smoothed vector exceeding this threshold matched the average sleep duration for the given work schedule. The segments of the smoothed vector above this threshold were classified as sleep, formulating average sleep patterns for each work schedule. Using this approach, we estimated the sleep history for a given seven-day work history by mapping the corresponding average sleep patterns to each work pattern (black arrow in Fig. 2b (iv)). This seven-day historical sleep pattern was repeated until the total estimated sleep history exceeded 300 days.

## Result

### Long-term sleep data is required to calculate the circadian phase that is robust to the subjective initial condition choice

Mathematical model simulation has been acknowledged as the most promising method for predicting DLMO (Reiter et al., 2020; Stone et al., 2020b), the gold standard marker of the circadian phase. Specifically, this mathematical model simulates the circadian rhythm of the core body temperature (CBT) and calculates DLMO by subtracting 7h from the minimal phase of CBT (CBTmin) (Hilaire et al., 2007; Stone et al., 2019; St Hilaire et al., 2020; Stone et al., 2020a; Huang et al., 2021; Lim et al., 2025). This method of simulating CBTmin utilizes light exposure information as an input of a mathematical model (Forger et al., 1999; Jewett et al., 1999; Hilaire et al., 2007). However, collecting accurate light exposure data is often challenging. One alternative approach is estimating light exposure profile from recorded sleep data: assuming light exposure during waking hours and no light during sleep (Postnova et al., 2016; Hong et al., 2021; Knock et al., 2021; Song et al., 2023; Varma et al., 2023; Lim et al., 2024; Song et al., 2024; Lim et al., 2025; Song et al., 2025) (Fig. 1a). This allows us to simulate the mathematical model and calculate CBTmin and DLMO from only the sleep-wake pattern data, which can be easily obtained from wearable devices.

Using sleep-wake pattern data, we can simulate the rhythm of the CBT after the initial condition (IC). While this IC affects subsequent CBT simulations and CBTmin calculations, these effects become negligible with sufficiently long-term sleep data because the simulated rhythms from different ICs eventually converge to the same phase in long-term simulations (i.e., entrainment occurs). In that convergent interval, CBTmin values become robust to IC (i.e. IC-robust CBTmin) and therefore reliable. However, the minimum length of the sleep data required for such IC-robustness remains unclear (Fig. 1b).

Here, we investigate the minimum sleep data length required to eliminate the effect of initial condition choice in CBTmin calculation, using sleep-wake pattern data from 28 shift workers collected via ActiWatch (mean = 56 days, 34-75 days). We first estimated light exposure profiles for each subject by assuming 250lux during awake and 0lux during sleep, utilizing this light exposure profile as an input for the mathematical model to simulate the CBT rhythm (Postnova et al., 2016; Hong et al., 2021; Knock et al., 2021; Song et al., 2023; Varma et al., 2023; Lim et al., 2024; Song et al., 2024; Lim et al., 2025; Song et al., 2025). This simulation was repeated with 24 distinct ICs, each representing 1:00, 2:00, …, and 24:00 on an entrained cycle obtained by a constant sleep pattern in 22:00-6:00 (See Methods for more details), resulting in 24 different trajectories of simulated CBTmin (Fig. 1c). We then evaluated the day on which the standard deviation (SD) of CBTmin values (excluding outliers; see Methods for more details) drops below 0.1h (i.e. convergence), and the impact of IC on CBTmin calculation becomes minimal (Ranga Suri et al., 2019).

The results showed that all 28 subjects required at least 18 days of data for CBTmin convergence, and about 50% of subjects needed at least 27 days to eliminate the effect of subjective IC choice in CBTmin calculation (Fig. 1c). Meanwhile, some subjects did not provide IC-robust CBTmin values even when we used their entire sleep data (>50 days). To identify the reason for this discrepancy, we compared the sleep patterns of three subjects who required 18 days (Fig. 1d (i)), 34 days (Fig. 1d (ii)), and >50 days (Fig. 1d (iii)) for reliable circadian phase estimation. The subject with the shortest duration (i.e., 18 days) maintained a consistent sleep pattern until the CBTmin trajectories from 24 ICs (colored dots ranging from red to blue in Fig. 1d (i)) converged. Meanwhile, another subject who required a longer duration (34 days) exhibited a more irregular sleep pattern (Fig. 1d (ii)). Even more irregular sleep patterns led to the failure of CBTmin trajectories to converge, even with 63 days of sleep data (Fig. 1d (iii)). These results suggest that subjects with irregular sleep patterns (e.g., shift workers) require longer-term sleep data for reliable circadian phase estimation. However, the data lengths used in previous studies for estimating the circadian phases of shift workers may have been inadequate. For instance, Huang et al. (Huang et al., 2021) used 7–14 days, and J.E. Stone et al. (Stone et al., 2019) used 5–21 days. Given our findings, these durations were likely too short to ensure that CBTmin values were independent of IC choice, raising concerns about the reliability of their circadian phase estimations.

### The effect of the initial condition can be reduced by estimating the sleep history

In real-world scenarios, as seen in previous studies (Stone et al., 2019; Huang et al., 2021), sleep-wake pattern data is usually of shorter duration than the minimal length needed to ensure the simulation’s independence from the choice of the IC (Fig. 2a (i)). In such cases, CBTmin calculations remain dependent on the subjective choice of IC. However, the IC is usually unknown in a real-world scenario, as it is determined by the circadian rhythm state at the initial time point (i.e., the beginning of the sleep data), which depends on the prior circadian rhythm, also unknown. Thus, previous studies have estimated the prior circadian rhythm and IC by reconstructing the sleep pattern preceding the available sleep data (i.e., sleep history; Fig. 2a (ii)) and simulating the mathematical model for this reconstructed history (Fig. 2a) (Stone et al., 2019; Huang et al., 2021)

To reconstruct the sleep history for IC decisions, two methods have been mainly used (Stone et al., 2019; Huang et al., 2021). One of the methods is the ‘regular sleep method’ (Huang et al., 2021), which assumes a constant sleep pattern at fixed times (e.g., 23:30-7:30) (Fig. 2b (i)). While this method is simple and requires no complex calculations, it fails to incorporate any personal sleep characteristics from recorded sleep data. To address this problem, a previous study proposed the ‘mean midsleep method’ (Stone et al., 2019), which determines the timing of constant sleep patterns based on personalized characteristics in recorded sleep data (Fig. 2b (ii)). Specifically, this method sets the historical sleep timing so that historical CBTmin aligns with the mean of midsleeps on days following non-night shifts (indicated bold in Fig. 2b (ii)), thereby preventing distortion of historical sleep patterns caused by the irregular work schedules of shift workers (Stone et al., 2019).

However, these two previous methods (regular sleep and mean midsleep methods) have considerable limitations. Their assumption of constant sleep fails to account for intra-individual variation (i.e., daily variation) in sleep history. This limitation manifests in shift workers with irregular sleep schedules as their daily sleep duration and timing drastically change. This may explain why previous studies show lower circadian phase prediction accuracy for shift workers compared to regular workers (Stone et al., 2019; Huang et al., 2021).

To overcome these limitations, we developed two new methods that consider individual sleep characteristics (Fig. 2b), including daily variation. First, to directly reflect the individual’s sleep pattern in the sleep history, we developed the ‘period-based sleep method’ (Fig. 2b (iii)), which assumes that a specific period from recorded sleep data has been constantly repeated in the sleep history. This repetition-based method has been adopted in the previous study (Mayer et al., 2023), yet the optimal segment of sleep data to repeat has remained unclear. Here, we propose a systematic method for selecting this repetition period. The starting point of the repetitional period was fixed on the first day of the recorded sleep data (Fig. 2b (iii)), while the endpoint was determined to connect seamlessly to the start point of the next repetitional period. Specifically, the endpoint was identified as a day before a day with a sleep pattern similar to that of the first day of recorded data. To find this day with a similar sleep pattern, we first identified the days with the same work patterns (indicated in bold in Fig. 2b (iii)) under the assumption that having the same work pattern increases the likelihood of having similar sleep patterns. For instance, if the first day involved off, all subsequent days with off were considered likely to have similar sleep patterns (indicated in bold in Fig. 2b (iii)). Among these, the day whose sleep pattern most closely resembled that of the first day was selected (Fig. 2b (iii); see Methods for more details). The day preceding this selected day was chosen as the endpoint of the repetitional period (dotted box in Fig. 2b (iii)). Finally, this period was repeated in the sleep history until the total sleep history’s length exceeded 300 days to ensure sufficient length for IC-robustness in the history’s simulation.

The period-based sleep method estimates sleep history using only recorded sleep data and its corresponding work data. However, there are cases where the work history is available even when the sleep history is unknown. For such cases, we developed the ‘work history-based sleep method’ (Fig. 2b (iv)), which estimates sleep history by utilizing the work history. In this method, we first calculated the average sleep patterns (box in Fig. 2b (iv)) for each work pattern by using the recorded sleep data and corresponding work data (See Methods for more details). We then reconstructed the sleep history with these average sleep patterns, particularly matching the average sleep with the work history (black arrow in Fig. 2b (iv)). For instance, if four days of an individual’s work history show three days of night shifts, followed by a day off before sleep data recording, the sleep history would be reconstructed with the average sleep pattern of night shifts applied to the first three days in the history, followed by the average sleep pattern of off applied to the next day. After obtaining seven days of sleep history based on the work history, we extended the history by repeating this seven-day sleep pattern until the length of the extended history exceeded 300 days to ensure sufficient length for IC-robustness in the history’s simulation.

### Our IC estimation methods provide more reliable circadian phase information than previous IC estimation methods

To compare which of the four methods (regular sleep, mean midsleep, period-based sleep, and work history-based sleep methods) is most suitable for obtaining reliable circadian phase, we used sleep data from 28 shift workers with lengths ranging from 34 to 75 days (average = 56 days; Fig. 3a). Using this real long-term sleep data (Fig. 3b (i)), we simulated 24 trajectories of CBTmin from 24 different ICs, as done in Fig. 1c. We determined the convergence point of these trajectories (SD < 0.1, excluding outliers; see Methods for more details) (Ranga Suri et al., 2019), and defined the CBTmin values at and after this convergence point as robust to ICs and therefore reliable. From this IC-robust range, we extracted short-term data (Fig. 3b (ii)) covering 20 days. Then, we used this short-term sleep data to estimate sleep history (Fig. 3b (iii)) and replaced the real sleep history with the estimated history. For this estimated sleep history, we simulated the mathematical model to obtain the IC (Fig. 3b (iv)). Using this IC, we recalculated the CBTmin for each of the 20 days in the short-term sleep data and compared them to the reliable CBTmin values obtained from the real sleep history by computing their difference (i.e., error). Then, for each day in the short-term sleep data (1^st^ – 20^th^), the percentage of cases where the error was less than 0.5h was calculated. This calculation was repeated for three mathematical models (simpler, higher-order, and nonphotic models).

**Fig. 3.**
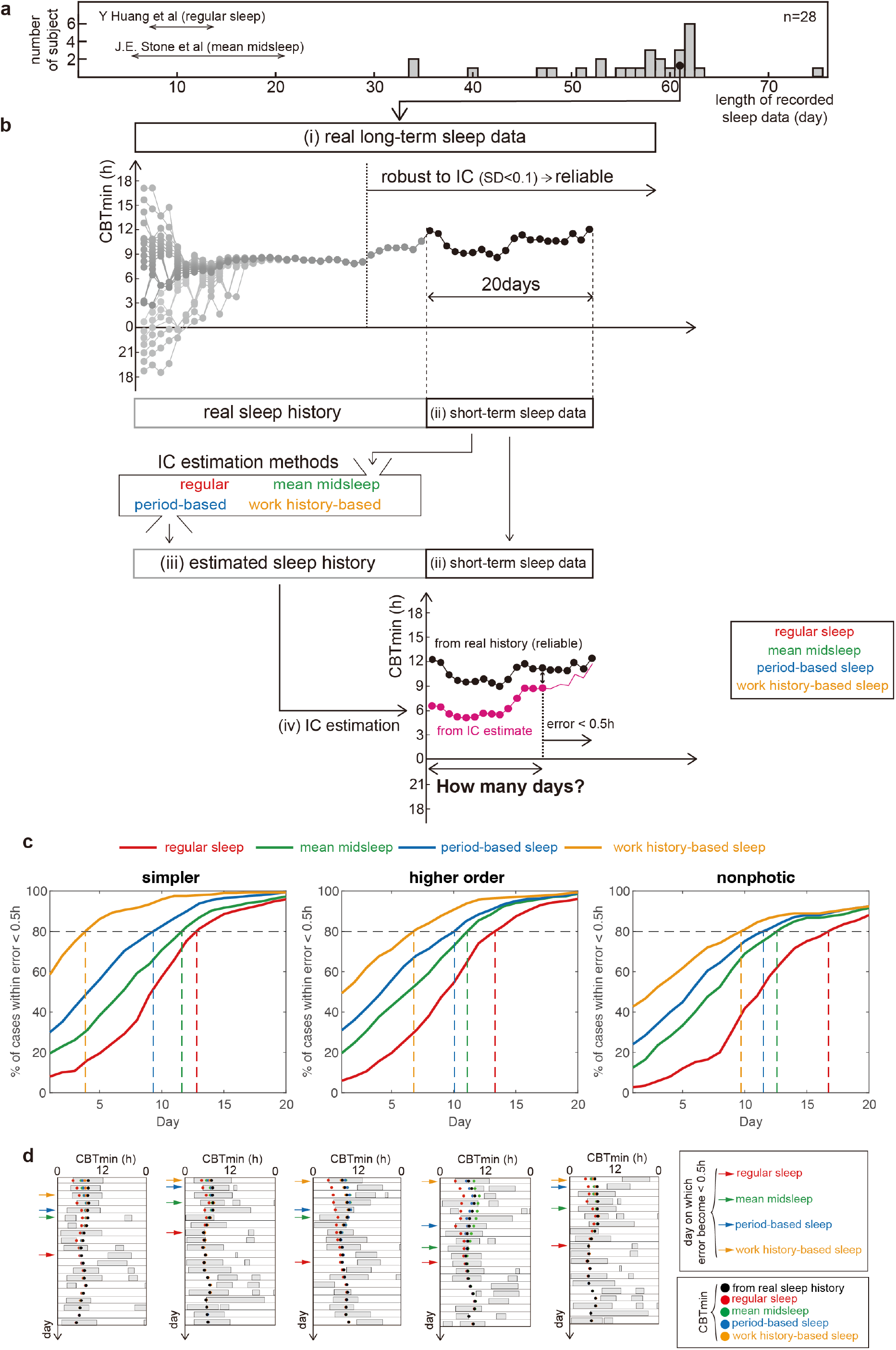
The period- and work history-based sleep methods provide performance superior to previous methods in estimating a reliable circadian phase similar to those obtained from real history. (a) To evaluate the performance of the four initial condition (IC) estimation methods (Fig. 2b), we used sleep data from 28 shift workers with an average data length of 56.04 days. (b) Using this real long-term sleep data (i) as input for the mathematical model, we simulated the CBT rhythm starting from 24 initial conditions as done in Fig. 1c. From this simulation, we identified the day on which 24 simulated CBTmin values converged, using the same definition of convergence described in Fig. 1c (i.e. SD<0.1, excluding outliers), and CBTmin values after this day were considered robust to IC, and therefore reliable. We examined how many days are required to estimate a reliable circadian phase when applying IC estimation methods. In particular, within the range in which CBTmin calculation was robust to IC, we extracted 20 days of short-term sleep data (ii). Using this short-term sleep data, we applied IC estimation methods and obtained the sleep history (iii) and IC value (iv). Then, the error between CBTmin values from the real history (i.e., reliable) and the IC estimate was computed on each day of the short-term data, and the days on which the error dropped below 0.5h were identified. (c) The plots showing the percentage of cases that succeeded in achieving an error below 0.5h each day (1^st^∼20^th^) across three mathematical model types: simpler, higher-order, and nonphotic models. 80% of cases achieved error<0.5h on about the 15^th^ and 13^th^ days on average in the regular sleep and mean midsleep methods, respectively. The day where 80% of cases achieve error<0.5h was moved forward when the period- and work history-based sleep methods were utilized, to about the 11^th^ and 7^th^ days in the period- and work history-based sleep methods, respectively. (d) Thus, when the same length of data is provided, the work history-based sleep method (orange) achieves reliable circadian phase estimation first, followed by the period-based sleep method (blue). On the other hand, using the previous methods, regular sleep (red) and mean midsleep (green), reach reliable estimation on later days. Hence, our IC estimation methods outperform previous approaches by providing reliable circadian phase estimation for a larger portion of days within the given sleep data.

When the previous methods, the regular sleep (red in Fig. 3c) and the mean midsleep (green in Fig. 3c) methods, were used to determine the IC, 80% of all cases achieved error<0.5h on approximately the 15^th^ and 13^th^ days, respectively, when averaged across the three model types (Fig. 3c). Meanwhile, with the period-(blue in Fig. 3c) and work history-based (orange in Fig. 3c) sleep methods, the day on which 80% of cases reached error < 0.5h was brought forward to the 11^th^ and 7^th^ days, respectively. This indicates that the period- and work history-based sleep methods provide 10 and 14 days of reliable circadian phase information, respectively, offering more circadian phase information than previous methods.

In summary, given the same duration of data, the circadian phases estimated using the work history-based sleep method (orange in Fig. 3d) converges to a reliable circadian phase first, followed by the period-based sleep method (blue in Fig. 3d), while the circadian phase estimates obtained by the previous approaches (red and green in Fig. 3d) converge to reliable circadian phases later than our methods. Hence, our approach allows a larger portion of circadian phase estimates from given sleep data to be considered reliable.

## Discussion

Although an IC decision is necessary for ODE-based circadian phase estimation, it remains unclear how IC uncertainty affects circadian phase estimation. In this study, utilizing sleep data from 28 shift workers collected via ActiWatch (mean length = 56 days, range = 34-75 days), we confirmed that IC robustness in circadian phase estimation is not ensured when sleep data is available for fewer than 18 days (Fig. 1). In other words, <18 days of sleep data, which is the typical data length used in previous studies (Stone et al., 2019; Huang et al., 2021), is insufficient to get reliable circadian phase information, unless the IC is accurate. To accurately estimate the IC, previous studies estimated the sleep history using the regular sleep and mean midsleep methods. These methods indeed reduced the days required for reliable circadian phase estimation from ≥18 days to 13 – 15 days on average in 80% of all cases (red and green in Fig. 3c). To further shorten this requirement, we developed two new approaches: the period- and sleep history-based sleep methods (Fig. 2b). Using our approaches, reliable circadian phase estimates can be achieved from only 11 and 7 days of sleep data, for the period- and work history-based sleep methods, respectively (blue and orange in Fig. 3c), on average in 80% of all cases. These results indicate that our study provides a solution to the IC problem in circadian phase estimation.

The superior performance of our approaches compared to classical methods (Fig. 3c) can be attributed to their ability to capture daily variations in sleep, thereby reconstructing a more realistic sleep history. Indeed, when we quantified the similarity between actual sleep histories and those generated by the four IC estimation methods (Text. S1 and Fig. S1), the work history–based method achieved the highest similarity, followed by the period-based, mean midsleep, and regular sleep methods—consistent with the results from CBTmin estimation (Fig. 3c). These findings suggest that future efforts to further improve the accuracy of sleep history estimation will, in turn, enhance the precision of IC construction and increase the reliability of CBTmin estimation.

Beyond the improvement in the accuracy of circadian phase estimation, our methods will also be beneficial for studies on clinical factors, such as mood, as they are closely linked to the circadian phase (McClung, 2011; Kim et al., 2017). For instance, by harnessing the daily information of circadian phases and mood states, Song et al. demonstrated that the circadian phase causes mood change (Song et al., 2024). In another study, Lim et al. proposed an algorithm that predicts the following day’s mood episodes in patients with mood disorders using today’s circadian phase (Lim et al., 2024). Given that these studies relied on daily circadian phase information, our IC estimation methods can enhance causal inferences from circadian rhythm to mood and improve the predictability of mood episodes based on the circadian phase by providing more accurate phase estimates.

Despite the advantages of our IC estimation methods, several limitations remain. One limitation is that our IC estimation method was validated in a relatively homogeneous sample. In particular, all participants were female, and they were nurses. Given that sleep patterns vary by gender (Jonasdottir et al., 2021) and occupation (Sun et al., 2015), the sleep patterns in our dataset may not fully represent the general population. To verify the general superiority of our methods, additional validation is required with males and with people in other occupations. In addition, four IC estimation methods were compared under a simplified light intensity assumption of 250 lux during wakefulness and 0 lux during sleep. As this assumption could influence the comparative performance of IC estimation methods, we repeated the analysis using a higher light intensity of 800 lux during wakefulness (Fig. S2). The results showed that the period- and work history–based sleep methods maintained their superiority over the regular and mean midsleep methods. These findings suggest that the superiority of our methods remains robust under more realistic light conditions, and further investigating this dependence will be an important direction for future research.

Another limitation is the challenge of calculating average sleep patterns in the work history-based sleep method. We calculated average sleep patterns for each of the subjects and each of their work schedules using their entire sleep data (average = 54 days). However, when only limited sleep data is available, a sleep pattern corresponding to a specific work schedule may be missing, making it impossible to calculate the average sleep pattern for that schedule. For instance, if only five days of sleep data are available for a work schedule of night-night-off-off-off, the average sleep pattern for an evening shift cannot be calculated. This limitation can be mitigated by estimating average sleep patterns for each work schedule through surveys, but it must be verified whether such an approach still maintains the highest accuracy for the work history-based sleep method. Additionally, further research is needed to determine how to calculate average sleep patterns for each work schedule when only short-term sleep data is available.

Although our method improves IC accuracy in circadian phase estimation, the overall accuracy is also influenced by the intrinsic error of the model (i.e., model error) in addition to IC error. For example, in the ODE models we used, the intrinsic period (τ) is assumed to be 24.2 hours for all individuals (Stone et al., 2019; Huang et al., 2021), while this value varies depending on an individual’s physiological state (Scheer et al., 2007; Stack et al., 2020; Skeldon et al., 2023). As a result, the ODE model may inaccurately estimate the circadian phase in some individuals, such as persons with lower intrinsic periods. Nevertheless, our IC estimation methods outperform classical methods regardless of the model type (Fig. 3c). Therefore, when a more advanced model is developed, integrating it with our IC estimation methods will enable even more accurate circadian phase estimation.

## Acknowledgments

This study was funded by grants from the Institute for Basic Science (IBS-R029-C3) (J.K.K.) and the Korean government (MSIT) (NRF-2019R1A2C1090643)(J.H.K.), and 2018 research award grants from the Korean Sleep Research Society (J.H.K.)

## Author Contributions

Data collection: J.H.K. Concept and design: D.L. and J.K.K. Development of the algorithm: D.L., J.K., and J.K.K. Analysis of data: D.L., J.K., and J.K.K. Drafting of the initial manuscript: D.L., J.K., and J.K.K. Critical revision of the manuscript for important intellectual content: D.L., J.K., and J.K.K.

## Conflict of Interest Statement

The authors have no potential conflicts of interest with respect to the research, authorship, and/or publication of this article.

## Data & Code Availability Statement

The sleep pattern data of 28 shift workers are protected and unavailable due to data privacy laws. Specific academic requests for access to these data should be directed to the authors (J.K.K.; J.H.K.). The code underlying this work will be made publicly available upon acceptance.

## Notes

### Competing Interest Statement

The authors have declared no competing interest.

